# Distinct eosinophil subsets are modulated by agonists of the commensal-metabolite and vitamin B3 receptor GPR109A during allergic-type inflammation

**DOI:** 10.1101/2022.08.04.502285

**Authors:** Rossana Azzoni, Kara J. Filbey, Rufus H. Daw, Maria Z. Krauss, Matthew R. Hepworth, Joanne E. Konkel, Edith M. Hessel, Yashaswini Kannan, John R. Grainger

## Abstract

Eosinophils are key contributors to allergic pathology, however, increasingly eosinophils are described to have important roles in organ health and immunoregulation. Factors that impact these diverse functions of eosinophils are not understood. Here we show in allergic-type lung inflammation, metabolically distinct populations of eosinophils can be identified based on expression of Siglec-F (Siglec-F^hi^ and Siglec-F^int^). Notably, the lung Siglec-F^hi^ population was responsive to the commensal microbiome, expressing the short-chain fatty acid receptor GPR109A. Animals deficient in GPR109A displayed augmented eosinophilia during allergy. Moreover, transferred GPR109A-deficient eosinophils released more eosinophil peroxidase than controls. Treatment with butyrate or vitamin B3, both GPR109A ligands, reduced Siglec-F^hi^ eosinophil frequency and activation, which was associated with apoptosis of Siglec-F^hi^ eosinophils. These findings identify GPR109A as an unappreciated regulator of glycolytic Siglec-F^hi^ eosinophils, raising the possibility of depleting pathological eosinophil populations in disease states while sparing those with homeostatic functions.

## INTRODUCTION

Eosinophils are a hallmark of type 2 immunity (Spencer and Weller, 2010) and are considered to be major mediators of pathology in allergic-type inflammation (Fahy, 2009; Lee et al., 2004).This is due to their capacity to release cytotoxic granule proteins, such as eosinophil peroxidase (EPX) as well as cytokines and lipid mediators, contributing to the ongoing inflammatory response and tissue damage (Liu et al., 2006; Moqbel and Odemuyiwa, 2008). However, in health eosinophils can perform important roles in regulating immune responses and homeostatic functions (Lee et al., 2010; Marichal et al., 2017). For example, eosinophils have been shown to maintain appropriate glucose metabolism and insulin levels in adipose tissue by producing IL-4 and subsequent M2 macrophage polarisation, which is necessary to maintain metabolic homeostasis (Wu et al., 2011).

One setting in which both inflammatory (Siglec-F^hi^) and homeostatic (regulatory) (Siglec-F^int^) eosinophil subsets have been identified is in the context of allergic-type lung inflammation (Mesnil et al., 2016; Abdala-Valencia et al., 2016). In this setting, Siglec-F^hi^ eosinophils were shown to be pro-inflammatory by enhancing the inflammatory response in the airways, while the Siglec-F^int^ population was found to be present in the healthy lung and suppress T cell proliferation by impacting dendritic cell (DC) activation (Mesnil et al., 2016).

While steroids are the first-choice therapy for allergic asthma, more recently biologic drugs targeting eosinophils globally, such as anti-IL-5, are being employed (Wechsler et al., 2012). Unlike tissue-resident macrophages, eosinophils are considered to have short half-lives and constantly replenished from bone marrow (BM). Given this, total systemic eosinophil depletion in disease settings may also limit delivery of eosinophils to organs where they have regulatory/homeostatic roles. Therefore, it is proposed that global depletion via biologics such as anti-IL-5 could result in unappreciated secondary complications. Understanding of eosinophil heterogeneity is still in its infancy, but identifying alternative approaches for targeting specific eosinophil subsets is of high therapeutic interest.

For other well-studied populations of immune cells, such as macrophages and T cells, differential functionality is often associated with alterations in metabolism (Viola et al., 2019; MacIver et al., 2013). In the case of eosinophils, it has been demonstrated that upon activation these cells enhance glycolysis but also mitochondrial respiration, both triggered by IL-5 and GM-CSF *in vitro* (Jones et al., 2020). However, whether different subsets of eosinophils *in vivo* are metabolically distinct during inflammation has not been well explored. If differential metabolism was established between subsets, this could help to inform candidate strategies to impact inflammatory populations over their regulatory or homeostatic counterparts.

One new avenue of therapies that target alternative pathways to steroids and anti-IL-5 in type 2 settings are commensal-derived metabolites, particularly short-chain fatty acids (SCFAs). These are a group of factors that are implicated in regulating type 2 immune responses (Dang and Marsland, 2019), including in allergic-type inflammation of the lung (Trompette et al., 2014; Thorburn et al., 2016). SCFAs have also been shown to target glycolytically active cells and to induce metabolic adaptations (Bachem et al., 2019). Much of the focus of these types of targets has been on their systemic effects, and although it has been suggested that commensal-derived factors can directly impact eosinophils *in vitro*, whether they are selective for different subsets is unknown.

Here we show that in the context of HDM-induced allergic-type inflammation, Siglec-F^hi^ eosinophils represent a metabolically distinct population to their Siglec-F^int^ counterparts. Lung Siglec-F^hi^ eosinophils, but not Siglec-F^int^ eosinophils, expressed high levels of the neutral amino acid transporter CD98, which correlated with greater activation of the mTOR signaling pathway and with enhanced metabolic activity. These features were induced in the lung environment but not in the circulation or bone marrow. Functionally, Siglec-F^hi^ eosinophils were pro-inflammatory and more activated than Siglec-F^int^ eosinophils. Siglec-F^hi^ eosinophils-expressing CD98 were not only present in allergic mice but are a common feature of lung inflammatory responses. Importantly, we found that Siglec-F^hi^ eosinophils were selectively impacted by oral butyrate treatment and had augmented expression of the SCFA receptor GPR109A. We found that intranasal administration of GPR109A agonists, such as niacin, could ameliorate HDM-induced lung eosinophilia and could spare systemic eosinophils, thus providing local benefit. These data highlight GPR109A as an unappreciated pathway important in regulating pathologic glycolytic Siglec-F^hi^ eosinophils and establish ligands of this pathway as candidate therapeutics for targeted treatment of allergic inflammation.

## METHODS

### Mice

Wild-type (WT) BALB/c and C57BL/6 female mice were purchased from Envigo and housed in individually ventilated cages under specific pathogen free (SPF) conditions. Germ-free (GF) and SPF controls were bred in-house and were on a C57BL/6 background. *Hcar2^−/−^* mice were on a C57BL/6 background (Tunaru et al., 2003) and were bred in-house and kindly shared by Prof. Stefan Offermanns. ΔdblGATA mice (Dyer et al, 2007) were on a C57BL/6 background and were bred in-house and kindly shared by Prof. Sheena Cruickshank. All experiments were approved by The University of Manchester Local Ethical Review Committee and were performed in accordance with the UK Home Office Animals (Scientific Procedures) Act 1986 and the GSK Policy on the Care, Welfare and Treatment of Animals.

### *In vivo* treatments and infections

#### House dust mite (HDM)

Mice were sensitised with 100 μg HDM extract (Greer Laboratories) in 40 μl Phosphate Buffer Saline (PBS; Sigma Aldrich) or with PBS alone given intranasally (i.n.) on day 0. Mice were challenged with 10 μg HDM extract in PBS or with PBS alone given i.n. on days 7-11. Animals were sacrificed 24 hours after the last HDM challenge (day 12).

#### Sodium butyrate and niacin

Sodium Butyrate (Sigma Aldrich) was administered in the drinking water at 300 mM for two weeks prior and during HDM treatment. In some experiments, sodium butyrate and niacin (Sigma Aldrich) were delivered i.n. in 40 μl of PBS at 30 mM twice a week for two weeks prior and during HDM treatment or during the challenge of HDM treatment alone.

##### Nippostrongylus brasiliensis

For infection with *N. brasiliensis*, 250 L3 larvae were injected subcutaneously (s.c.) on day 0 and tissues were harvested on day 6. *N. brasiliensis* larvae were cultured as described previously (Lawrence et al., 1996).

##### Schistosoma mansoni

For infection with *S. mansoni*, 20 cercariae were injected s.c. on day 0 and 7 and subsequently 80 cercariae were injected on day 14. Tissues were harvested on day 28. *Biomphalaria glabrata* snails exposed to *S. mansoni* miracidia (NMRI strain) were acquired from Biodefense and Emerging Infections research repository or from Karl Hoffmann (Aberystwyth University) and isolated as described previously (MacDonald et al., 2001).

### Influenza

For infection with Influenza virus (PR8 strain), mice were administered 30 μl of virus suspension (5 pfu) on day 0 and tissues were harvested on day 3 or day 7.

### Euthanasia and tissue harvesting

In all experiments, mice were euthanized by exposure to a rising concentration of CO2 and tissues harvested at the indicated times post-treatment.

### Murine tissue preparation and cell isolation

#### Lung

Tissues were chopped finely in 100 μg/ml liberase TL (Roche) and 250 μg/ml DNaseI (Sigma Aldrich) and digested for 20 minutes at 37°C in a shaking incubator. 5 mM EDTA (Sigma Aldrich) was added for 5 minutes, after which the resulting suspension was then passed through a Corning^®^ 70 μm cell strainer (Sigma Aldrich). After pelleting by centrifugation (500 x g, 5 minutes, 4°C), the pellet was resuspended in ACK lysing buffer (Lonza) for 3 minutes on ice. Suspensions were then washed with PBS and resuspended in complete RPMI 1640 (Sigma Aldrich) (supplemented with 10 mM HEPES (Sigma Aldrich), 5 μg/ml penicillin-streptomycin (Sigma Aldrich), 2 mM L-glutamine (Sigma Aldrich), Minimal nonessential amino acids 100x (Sigma Aldrich)) containing 10% foetal calf serum (FCS; Sigma Aldrich), until staining.

#### Bronchoalveolar lavage (BAL)

BAL was performed three times with 400 μl PBS containing 5mM EDTA and 2% FCS. Fluid was centrifuged and supernatant was collected and stored at −80°C for cytokine detection. Pellets were resuspended in complete RPMI containing 10% FCS until staining.

#### Blood

Blood was collected into EDTA-coated syringes by cardiac puncture. Suspensions were washed and resuspended twice in ACK lysing buffer for 3 minutes on ice. Suspensions were then washed with PBS and resuspended in complete RPMI containing 10% FCS until staining.

#### Bone-marrow

Femurs were collected and ends removed. Each bone was placed in 0.5 ml Eppendorf tubes pierced with a 21G syringe. These in turn were inserted into 1.5 ml Eppendorf tubes and pulsed to maximum speed in a microcentrifuge. Cell pellets were resuspended in ACK lysing buffer for 3 minutes on ice. Suspensions were then washed with PBS and resuspended in complete RPMI containing 10% FCS until staining.

### Flow cytometry

#### Surface marker staining

Single-cell suspensions (5 × 10^5^–2 × 10^6^ total cells) were washed with PBS and stained with the Live/Dead Fixable UV dead cell stain kit (ThermoFisher) to exclude dead cells. Subsequently, cells were stained in the dark for 20 minutes at 4°C with fluorochrome- or biotin-conjugated antibodies in PBS containing anti-CD16/CD32 (2.4G2; BioXcell) in the dark. Cells were washed and, where necessary, incubated for a further 15 minutes with fluorochrome-conjugated streptavidin and then washed. In some cases, cells were immediately acquired live, or alternatively, after further washing, cells were fixed in 2% paraformaldehyde (PFA; Sigma Aldrich) for 10 minutes at room temperature and ultimately resuspended in PBS before acquisition.

Cells were stained with: CD11b (M1/70), CD11c (N418), CD45 (30F11), CD98 (RL388), CD125 (DIH37), Siglec-F (E50-2440) from BD and CD101 (REA301) from Miltenyi. The lineage antibody cocktail for excluding lymphocytes, other granulocytes and monocytes/macrophages for eosinophil analysis included TCRβ (H57-597), CD3 (17A2), B220 (RA3-6B2), Ly6G (1A8), Ly6C (HK1.4), CD19 (1D3/CD19), NK1.1 (PK136) and Ter119 (TER-119) from BioLegend.

#### Annexin V and 7-AAD staining

Mixed BM cultures were incubated with IL-33, GM-CSF and IL-5 (Peprotech) in the presence or absence of niacin for 24 hours. Single-cell suspensions (5 × 10^5^ - 2 × 10^6^ total cells) were washed with PBS and resuspended in 1X Annexin binding buffer (Thermofisher) and washed once. Each sample was then incubated with 2.5 μl of Annexin V for 15 minutes at room temperature in the dark. Cells were washed and resuspended in Annexin binding buffer. Each sample was then incubated with 2.5 μl of 7-AAD for 15 minutes at room temperature in the dark. Cells were immediately acquired live.

#### Intracellular staining

To assess protein phosphorylation, cells were fixed and permeabilised using BD Phosphoflow kit according to manufacturer instructions. Cells were stained with p-mTOR (MRRBY) and pS6 (cupk43k) from ThermoFisher.

#### Sample acquisition and analysis

Appropriate compensation was performed using UltraComp eBeads^™^ (Invitrogen) with single colour stains and samples were acquired on a BD LSRFortessa^™^ running BD FACSDiva 8 software (BD) and exported as FCS files. Data was analysed using FlowJo software (TreeStar) and gates were set using fluorescence-minus-one controls.

#### Fluorescence activated cell sorting (FACS)

To sort eosinophil populations from the lung (L/D^−^Lin^−^CD11b^int/hi^Siglec-F^int/hi^) single-cell suspensions were prepared and resuspended in PBS with 2% FCS and 2 mM EDTA. Sorting was performed using a FACSAria Fusion (BD) and sorted cells were collected in complete RPMI with 20% FCS and stored on ice for use in cytospins and *in vitro* cell culture assays.

#### Eosinophil isolation and transfer into ΔdblGATA mice

Bone-marrow eosinophils were isolated using EasySep^™^ Mouse Biotin Positive Selection Kit II (Stem Cell Technologies) according to manufacturer instructions. Cells were resuspended in PBS and 3×10^5^/50 μl were intranasally administered to ΔdblGATA mice on day 10 of the HDM challenge model.

### Histology

#### Cytospin preparation and staining

Sorted lung or BM eosinophils were mounted on SuperFrost Plus^™^ Adhesion Slides (Thermofisher) using a Cytospin centrifuge (Cytospin 4; Thermo Fisher Scientific) operating for 5 minutes at 500 rpm. Cells were fixed with ice-cold methanol and stored at room temperature. Unspecific antibody binding on sorted cells was blocked with TSA blocking reagent (PerkinElmer) for 30 minutes. Sections were then incubated with either mouse anti-rabbit GPR109A (1:40; Abcam), mouse anti-rabbit GPR41 (1:50; Invitrogen) or Cy3-conjugated mouse anti-rabbit GPR43 (1:50; Bioss) for 1 hour at room temperature. Primary antibody staining was followed by secondary antibody staining with FITC goat anti-rabbit (1:200; Invitrogen) for 1 hour at room temperature. Cells were mounted with ProLong^™^ Gold Antifade Mountant with DAPI (Thermofisher).

#### Image acquisition

Fluorescent images were collected on a Zeiss Axioimager.D2 upright microscope using either a 10x, 20x or 40x objectives and captured using a Photometrics Coolsnap HQ2 camera through Micromanager software v. 1.4.23.

All images were processed and analysed using Fiji Image J (http://imagej.net/Fiji/Downloads).

### Quantitative polymerase chain reaction (PCR)

#### RNA and DNA extraction

Tissues were collected into RNAlater^™^ stabilisation solution (Invitrogen) and stored at −80°C. To isolate RNA, 1 ml TRIzol^™^ reagent (Invitrogen) was added to tissues transferred into Lysing Matrix D tube (MP Biomedicals). Tissues were homogenised using a MP Biomedicals FastPrep-24^™^ Classic Grinder (4.0 m/s, 40 seconds) and rested on ice. Tissue suspensions were transferred to a clean Eppendorf tube and 200 μl chloroform (Sigma Aldrich) was added. Tubes were manually shaken for 15 seconds and rested at room temperature for 5 minutes. Tubes were centrifuged (12,000 x g, 15 minutes, 4°C) and the upper aqueous layer was transferred into a clean Eppendorf tube. To precipitate RNA, 500 μl propan-2-ol (Sigma Aldrich) was added and samples were incubated at room temperature for 10 minutes. Tubes were centrifuged (12,000 x g, 10 min, 4°C) to pellet the RNA. The supernatant was removed by tipping, and the RNA pellet was washed by vortexing with 1 ml 75% ethanol. Samples were centrifuged once more and RNA pellets were air-dried for 30-60 minutes at room temperature before resuspending in 30 μl RNase-free water (Qiagen). The quality and quantity of RNA was measured using a NanoDrop Microvolume spectrophotometer. Samples with 260/280 values of 1.60-2.00 and 260/230 values of 1.50-2.20 were considered of good quality. Samples were stored at −20°C until cDNA synthesis.

#### cDNA synthesis

RNA concentrations were normalised across samples through dilution with RNase-free water. cDNA synthesis was carried out using High-Capacity cDNA Reverse Transcription Kit (Applied Biosystems) according to manufacturer’s instructions and a Veriti 96-well thermal cycler.

#### Quantitative PCR (qPCR) and analysis

cDNA samples were diluted 1:8 in nuclease free water. Each sample was added in triplicate per RNA transcript on a 384 well qPCR plate (Applied Biosystems). Each well contained 4.5 μl cDNA, 5 μl Fast SYBR^™^ Green Mastermix (Applied Biosystems) and 0.5 μl 10 μM forward and reverse primers (Table 1). Samples were amplified using a QuantStudio 12 K Flex system.

**Table 1:**
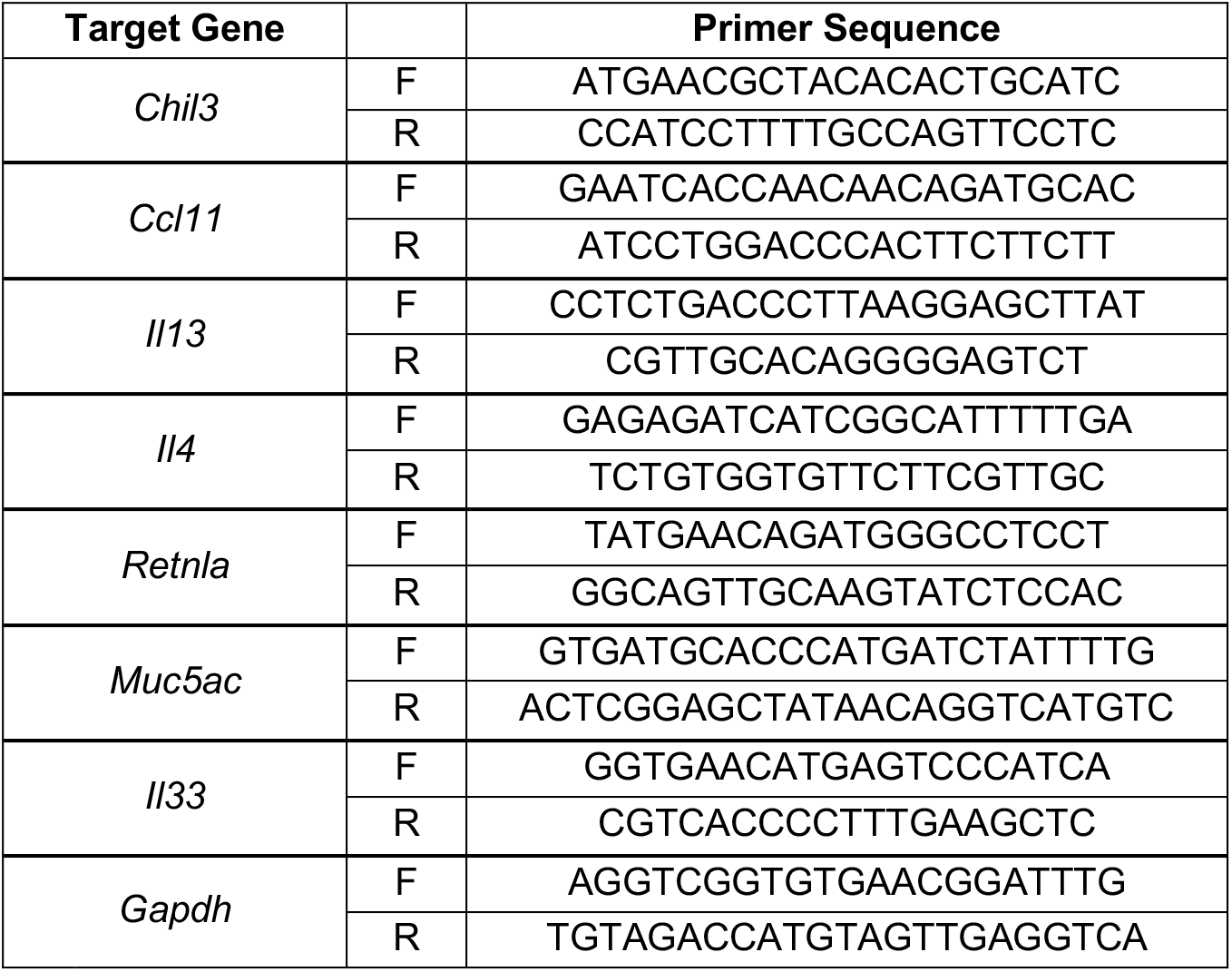
PCR primer sequences

Relative expression of target genes within experimental treatment groups was determined through comparison to an internal control gene (*Gapdh*) and the experimental control group.

### Supernatant analysis

#### Enzyme-linked immunosorbent assay (ELISA)

Supernatants of cultured eosinophils were screened for IL-6, IL-4 and EPX (Table 2). Appropriate primary antibodies were diluted in ELISA coating buffer (Biolegend) and added to 96 well high binding ELISA plates (Greiner Bio-One) overnight at 4°C. The coating solution was then flicked off and the plate was washed three times with PBS containing 0.1% Tween^®^ 20 (Sigma Aldrich). Plates were then blocked with PBS containing 10% FCS for 1 hour at room temperature. The blocking solution was flicked off and samples and standards were added and incubated overnight at 4°C. Plates were washed three times and secondary antibodies were then added and incubated for 2 hours at room temperature. Following washing for three times, streptavidin-peroxidase was added and incubated for 30 minutes at 37°C. The plate was finally washed eight times and was developed with the TMB substrate kit (Biolegend) according the manufacturer’s instructions. The reaction was stopped by adding Stop Solution for TMB substrate (Biolegend). The plates were read using a Tecan Infinite M200 PRO microplate reader running I-control 1.9 software at 450 nm, with reference of 570 nm subtracted.

**Table 2:**
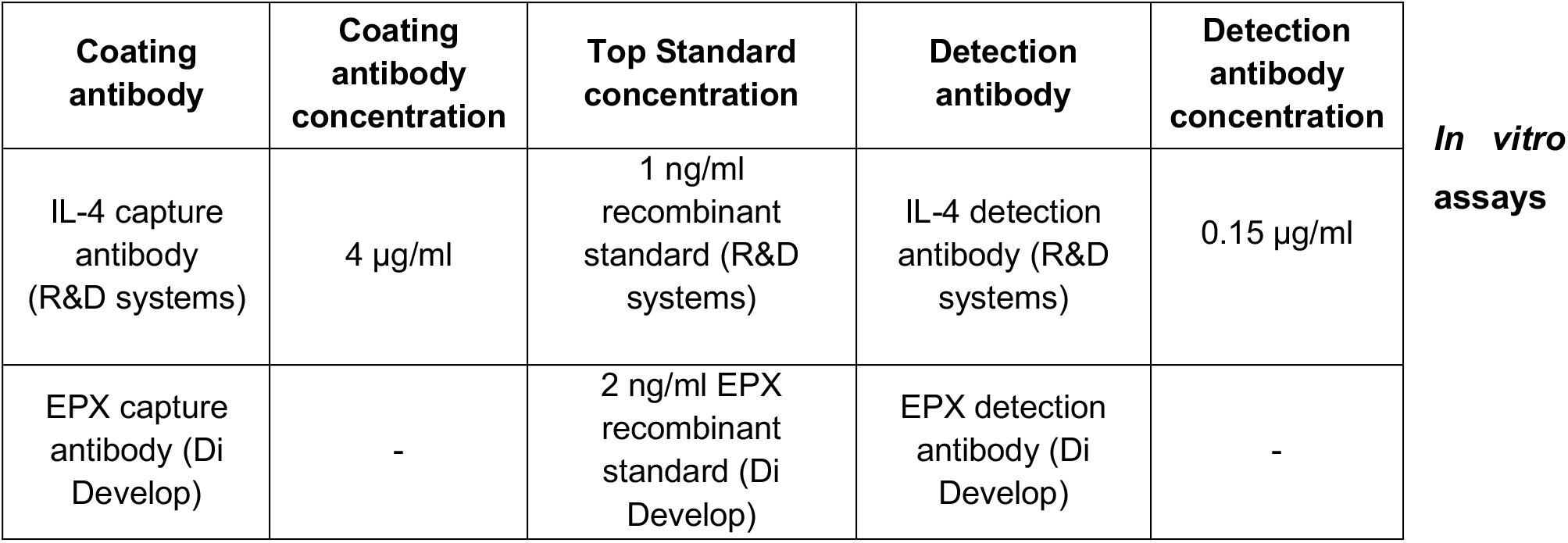
Antibodies and standard concentrations used for ELISA.

#### Seahorse extracellular flux analysis

Sorted eosinophils were plated at 150,000 cells/well and allowed to adhere for at least 1 hour. ECAR and OCR were measured in XF media (modified DMEM containing 2 mM L-glutamine) under basal conditions and in response to 10 mM glucose, 2.5 μM oligomycin, 100 mM 2-DG using a 96-well extracellular flux analyzer XFe-96 (Seahorse Bioscience).

#### Eosinophil stimulation

Sorted lung eosinophils were resuspended in complete RPMI with 10% FCS and added to a 96 well plate at a density of 1×10^6^ cells/ml. A23187 (Sigma Aldrich) was added at 5 μM and samples were incubated for 3 hours at 37°C. Cell supernatants were collected and analysed by ELISA.

#### Statistics

Statistical analyses were performed using Prism (8.0; GraphPad software) and data were presented as mean ± SEM in all cases. Two experimental groups were compared using a Student’s t test for unpaired data. Where more than two groups were compared, a one-way ANOVA or two-way ANOVA with Bonferroni’s correction was used. Significance was set at P ≤ 0.05.

## RESULTS

### Metabolically distinct Siglec-F^hi^ and Siglec-F^low^ eosinophils are a feature of allergic-type inflammation

To determine the mechanisms regulating distinct eosinophil populations during allergic inflammation, we employed a mouse model of house dust mite (HDM) allergic asthma (Hammad et al. 2010) (**Supplementary Fig. 1 A**). Eosinophils were initially identified based on their Siglec-F and CD11b expression. Alveolar macrophages (AMs) could be excluded as although Siglec-F^+^ they lack CD11b expression (**Supplementary Fig. 1 B**). As reported previously (Mesnil et al., 2016), in the healthy lung tissue a single population of Siglec-F^int^CD11b^int^ eosinophils was detectable by flow cytometry (**Fig. 1 A – C**). This population was also detected at extremely low levels in the naïve bronchoalveolar lavage (BAL) (**Fig. 1 A – C**). Contrasting this, during HDM-mediated allergic inflammation, total eosinophil numbers increased (**Fig. 1 B**) that were dominated by SiglecF^hi^CD11b^hi^ eosinophils in both the lung tissue and the BAL (**Fig. 1 A**).

**Figure 1:**
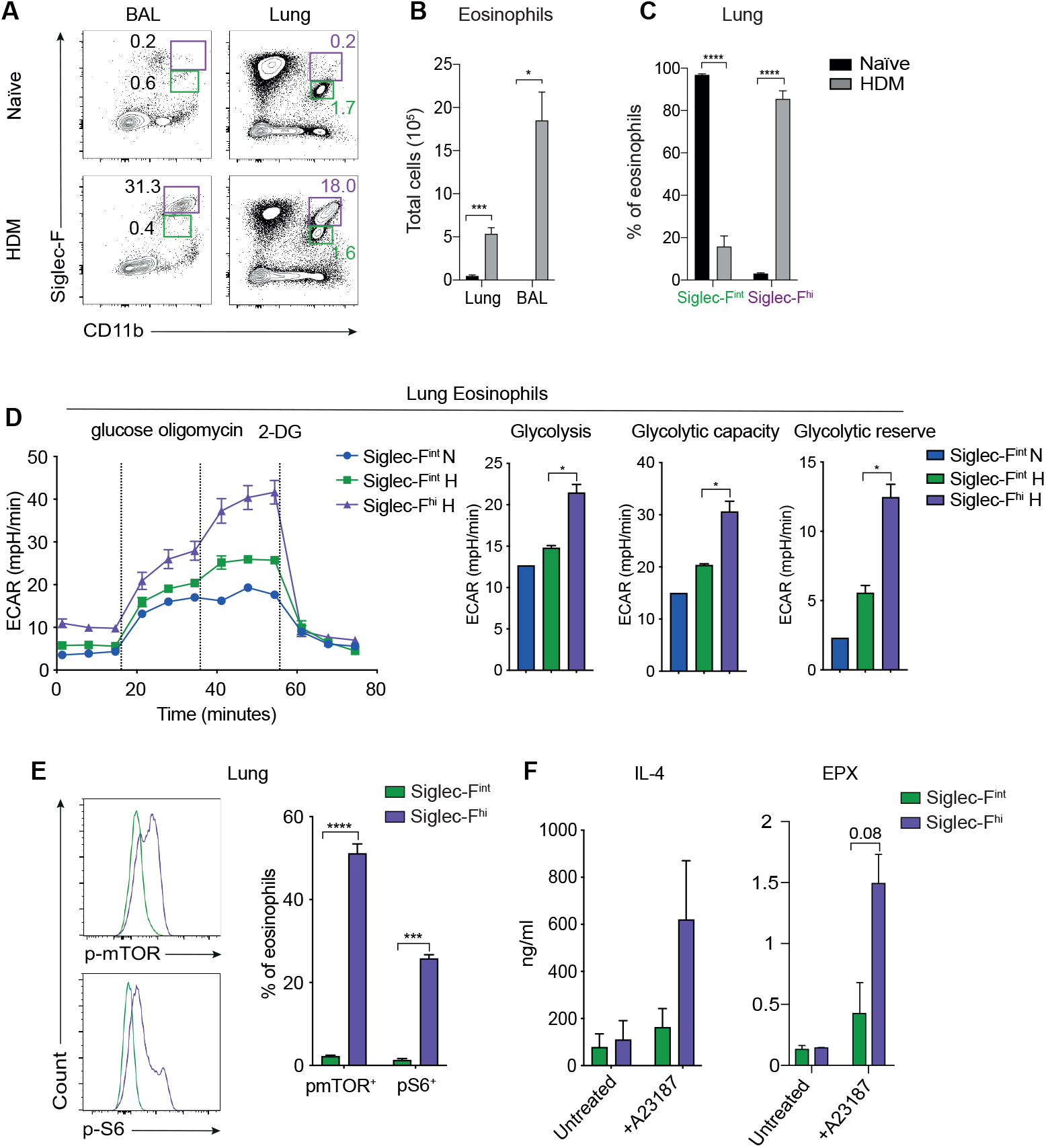
Metabolically distinct subsets of eosinophils are present in the allergic lung. **(A and B)** Eosinophils were identified by flow cytometry in the lung and BAL of naïve and HDM-treated mice. **(A)** Representative flow cytometry plot quantifying Siglec-F^hi^ (Purple gate) and Siglec-F^int^ (Green gate) eosinophils. **(B)** Total eosinophils in lung and BAL. **(C)** Frequency of Siglec-F^int^ and Siglec-F^hi^ eosinophils of total eosinophils. **(D)** ECAR of sorted lung eosinophils (Siglec-F^hi^CD11b^hi^ and Siglec-F^int^CD11b^int^) from naïve and HDM-treated mice at baseline and after sequential treatment (vertical lines) with glucose, oligomycin or 2-DG to measure glycolysis, glycolytic reserve and glycolytic capacity. **(E)** Frequency of Siglec-F^int^ and Siglec-F^hi^ eosinophils expressing p-mTOR and p-S6 quantified directly ex vivo by flow cytometry. **(F)** IL-4 and EPX levels were determined by ELISA in the supernatant from sorted lung Siglec-F^int^ and Siglec-F^hi^ eosinophils cultured in the presence or absence of A23187. Error bars show ± SEM. n = 2-6 per experiment. Data are representative of 3 independent experiments. Statistical comparisons were performed with Student’s t test (B, C and F) or one-way ANOVA (D): p < 0.05; **p < 0.01; ***p < 0.001.

Siglec-F^hi^ eosinophils are thought to be pathologic in the context of lung allergy, while Siglec-F^int^ eosinophils have been implicated in immunoregulation by limiting DC activation and reducing Th2 responses (Mesnil et al., 2016). Although not demonstrated for Siglec-F^hi^ and Siglec-F^int^ eosinophils, differential metabolism is associated with inflammatory versus immunoregulatory functions in other myeloid populations (Viola et al., 2019; Sadiku et al., 2021; Zuo and Wan, 2020). Thus, we investigated whether Siglec-F^hi^ eosinophils had a distinct metabolic potential to their Siglec-F^int^ counterparts. We performed a glycolytic stress test on sorted Siglec-F^hi^ and Siglec-F^int^ lung eosinophils from HDM-treated mice alongside Siglec-F^int^ eosinophils from naïve animals (**Fig. 1 D**). The assay analyses the rate of glucose catabolism into lactate by measuring extracellular acidification rates (ECAR) (Mookerjee et al., 2016). Cells were firstly incubated with glucose to trigger glycolysis and subsequently with oligomycin to inhibit mitochondrial ATP synthase, further driving glycolysis-mediated ATP production. Finally, the eosinophil populations were treated with 2-deoxy-D-glucose (2-DG), which inhibits glycolysis (Mookerjee et al., 2016). This approach allowed measurement of glycolysis (conversion of glucose to lactate), glycolytic capacity (maximum rate of glycolysis) and glycolytic reserve (difference between glycolytic capacity and glycolysis) (Mookerjee et al., 2016) (**Fig. 1 D**). Siglec-F^hi^ eosinophils from the HDM-treated group demonstrated greatly increased glycolysis, glycolytic capacity and glycolytic reserve compared to Siglec-F^int^ eosinophil populations from naïve and HDM-treated animals (**Fig. 1 D**).

Correlating with enhanced glycolysis, intracellular phospho-staining of Siglec-F^hi^ eosinophils also revealed that mammalian target of rapamycin (mTOR) and ribosomal protein S6 were phosphorylated, suggesting augmented nutrient uptake (Yerlikaya et al., 2016) compared to Siglec-F^int^ eosinophils (**Fig. 1 E**). Additionally, as expected, given their more glycolytic metabolic state (Soto-Heredero et al., 2020) Siglec-F^hi^ eosinophils released more IL-4 and eosinophil peroxidase (EPX) upon global activation with the Ca^2+^ ionophore A23187 (**Fig. 1 F**).

Taken together these findings demonstrate that two metabolically distinct populations of eosinophils are present in the lung environment during allergic-type lung inflammation.

### Specific characteristics of glycolytic Siglec-F^hi^ eosinophils are present in the lung environment

We further investigated the phenotype of the glycolytic Siglec-F^hi^ eosinophils compared to Siglec-F^int^ eosinophils present in the lung. To this end, we screened a set of markers by flow cytometry that have been associated with myeloid cell, or eosinophil function and activation specifically. As reported (Mesnil et al., 2016), Siglec-F^hi^ eosinophils expressed high levels of the T cell proliferation suppressor CD101 (Schey et al., 2016) along with the integrin CD11c (Abdala-Valencia et al, 2015), the inhibitory receptor CD300c/d (Rozenberg et al., 2018) and the early activation marker CD69 (Wang et al., 2004) (**Fig. 2 A**). They were also found to express higher levels of the type 2 effector molecule RELM-α (Sutherland et al., 2018) and the proliferation marker Ki-67 (suggestive of recent recruitment from the bone marrow (BM), compared to the Siglec-F^int^ population (**Fig. 2 A**). Of note, Siglec-F^hi^ eosinophils expressed the large amino acid transporter CD98 (LAT1) that facilitates functional transport of neutral amino acids (Cantor and Ginsberg, 2012), again supporting their enhanced metabolic activation (**Fig. 2 A**). Indeed, all markers investigated had greater expression on Siglec-F^hi^ eosinophils, except the co-stimulatory molecule CD86 (Woerly et al., 1999), compared to their Siglec-F^int^ counterparts (**Fig. 2 A**).

**Figure 2:**
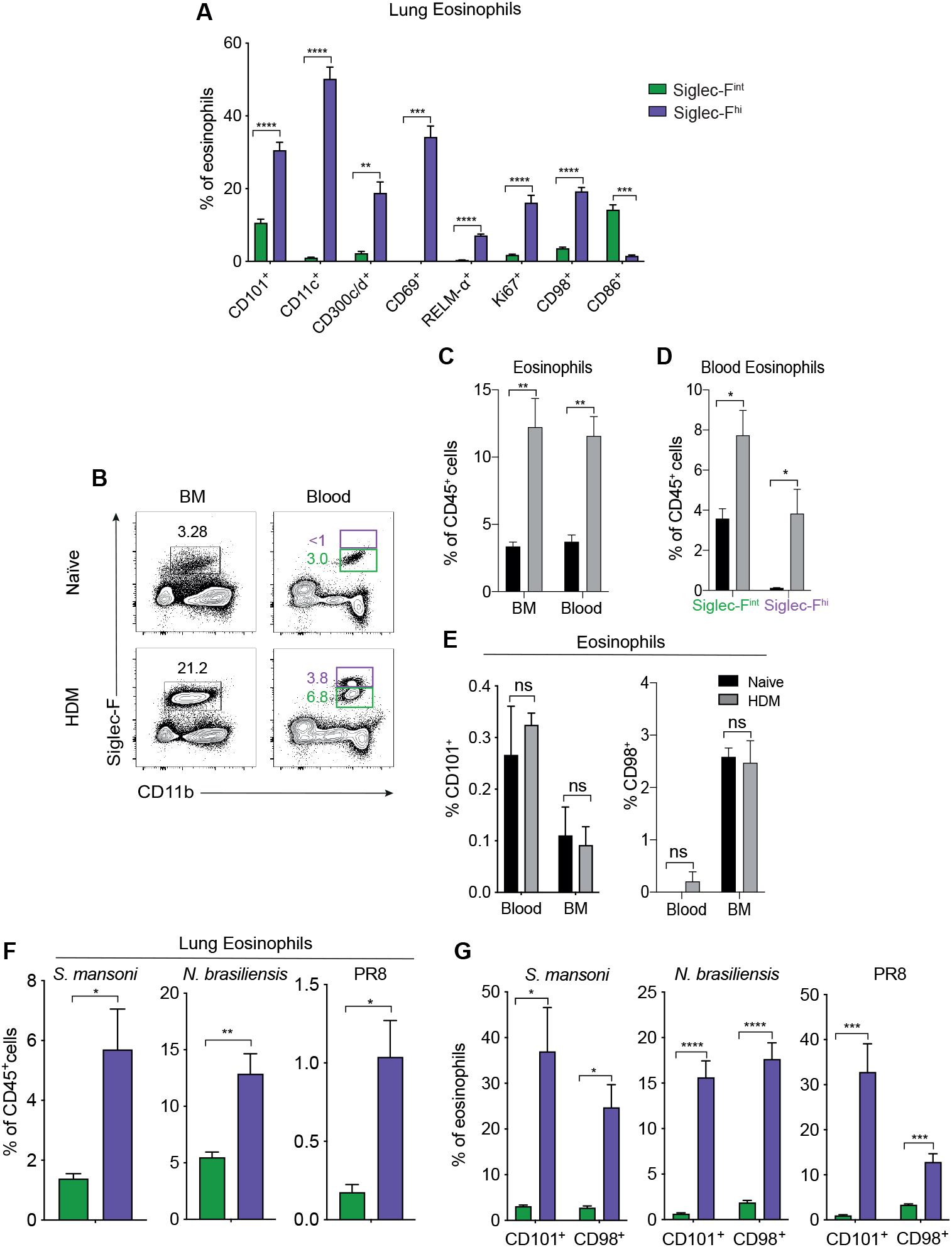
Siglec-F^hi^ eosinophils acquire specific characteristics in the lung during inflammation. **(A)** Frequency of Siglec-F^hi^ and Siglec-F^int^ eosinophils expressing CD101, CD11c, CD300c/d, CD69, RELM-α, Ki67, CD98 and CD86 was quantified by flow cytometry in the lung of HDM-treated mice. **(B and C)** Eosinophils were quantified by flow cytometry in the BM and blood of naïve and HDM-treated mice and expressed as a frequency of CD45^+^ cells. **(B)** Representative flow cytometry plot quantifying Siglec-F^hi^ (Purple gate) and Siglec-F^int^ (Green gate) eosinophils. **(C)** Frequency of eosinophils within CD45^+^ population in BM and Blood. **(D)** Frequency of Siglec-F^int^ and Siglec-F^hi^ eosinophils of total eosinophils. **(E)** Frequency of eosinophils expressing CD101 and CD98 quantified by flow cytometry. **(F)** Eosinophils were quantified by flow cytometry and expressed as a percentage of CD45^+^ cells in the lungs of naïve mice (green) or those infected with *N. brasiliensis*, *S. mansoni* or the PR8 strain of influenza (purple). **(G)** Frequency of eosinophils expressing CD98 and CD101 were quantified by flow cytometry in the lungs of naïve (green) and mice infected with *N. brasiliensis*, *S. mansoni* and PR8 strain of influenza (purple). Error bars show ± SEM. n = 3-6 per experiment. Data are representative of 3 independent experiments. Statistical comparisons were performed with Student’s t test: p < 0.05; **p < 0.01; ***p < 0.001.

During allergy, eosinophil production is augmented in the BM leading to subsequent increases in circulating eosinophils (Fahy, 2009; Inman 2000). Siglec-F^hi^ eosinophils have been reported in the circulation (Mesnil et al., 2016). As expected, we observed these described alterations to the BM and blood eosinophil compartment in our model (**Fig. 2 B – D**) Interestingly, circulating Siglec-F^hi^ eosinophils did not express CD101 or CD98, suggesting that they take on distinct functional and metabolic changes only in the lung environment (**Fig. 2 E**).

The presence of Siglec-F^hi^CD101^+^CD98^+^ eosinophils in lung-tissue was not restricted to HDM lung allergy, as it was a common feature of lung inflammation, including during Th2-driving parasitic *Nippostrongylus brasiliensis* and *Schistosoma mansoni* infection (**Fig. 2 F and G**) (**Supplementary Fig. 2 A and B**). They were also present, at lower frequency, during infection with the PR8 strain of influenza that drives a strong type I IFN response (**Fig. 2 F and G**) (**Supplementary Fig. 2 C**).

### Siglec-F^hi^ glycolytic eosinophils are modified by systemic exposure to the commensal-derived metabolite butyrate

Microbial-derived metabolites, such as SCFAs, are important regulators of pro-inflammatory responses (Maslowski et al., 2009; Trompette et al, 2014; Cait et al., 2018) and particularly glycolytic populations (Schulthess et al., 2019). Moreover, the SCFA butyrate has been reported to regulate eosinophil survival via its action as a histone deacetylase (HDAC) inhibitor (Theiler et al., 2019). We therefore investigated whether administration of butyrate could represent an unappreciated mechanism to selectively target the more glycolytic Siglec-F^hi^ eosinophil population.

Butyrate was administered in the drinking water of BALB/c mice for a 2-week period prior to and then during HDM sensitisation and exposure (**Supplementary Fig. 3 A**). Allergic mice treated with butyrate demonstrated reduced frequencies of Siglec-F^hi^ eosinophils in the lung compared to controls (**Fig. 3 A – C**). This was specific to the Siglec-F^hi^ subset as frequencies of Siglec-F^int^ eosinophils were unaffected by butyrate treatment (**Fig. 3 C**). Orally administered butyrate had systemic effects on eosinophils with reduced frequencies of eosinophils evident in the BM (**Fig. 3 A and B**). The phenotype and activation of Siglec-F^hi^ eosinophils in the lung were also impacted with reduced expression of both CD98 and CD101 (**Fig. 3 D**). Broader effects on the type 2 response were also apparent in butyrate-treated animals, including reduced gene expression of Th2-associated cytokines (IL-4 and IL-13) compared to HDM alone, as well as reduced type 2 effector molecules such as RELM-α and Ym-1 (**Supplementary Fig. 3 B**). These findings demonstrate that systemic alterations to commensal-derived SCFA can modify the Siglec-F^hi^ eosinophil population.

**Figure 3:**
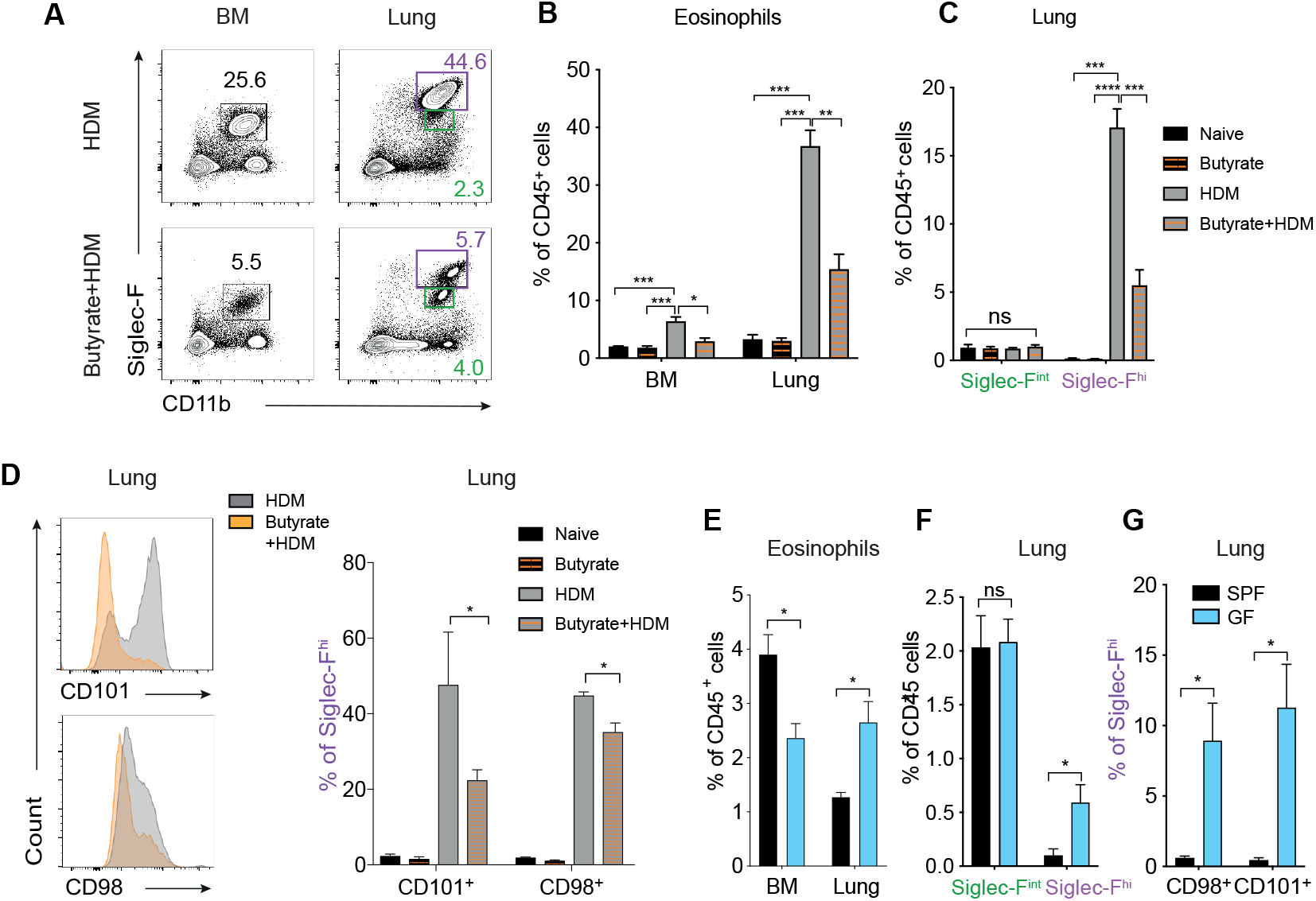
Siglec-F^hi^ eosinophils are modified by commensal-derived metabolites. **(A – C)** Eosinophils were quantified by flow cytometry in the lung and BM of naïve mice, mice treated with butyrate alone, mice treated with HDM alone and mice treated with HDM and butyrate. **(A)** Representative flow cytometry plot quantifying Siglec-F^hi^ (Purple gate) and Siglec-F^int^ (Green gate) eosinophils. **(B)** Frequency of eosinophils within CD45^+^ population in BM and Lung. **(C)** Frequency of Siglec-F^int^ and Siglec-F^hi^ eosinophils within the CD45^+^ population in the lung. **(D)** The percentage of Siglec-F^hi^ eosinophils expressing CD101 and CD98 was quantified by flow cytometry in the lung of naïve mice, mice treated with butyrate alone, mice treated with HDM alone and mice treated with HDM and butyrate. **(E)** Eosinophils were quantified by flow cytometry in the BM and lung of SPF and GF mice and expressed as a frequency of CD45^+^ cells. **(F)** Frequency of Siglec-F^int^ and Siglec-F^hi^ eosinophils of total CD45^+^ cells in the lung. **(G)** The frequency of Siglec-F^hi^ eosinophils expressing CD101 and CD98 was quantified by flow cytometry in the lung of SPF and GF mice. Error bars show ± SEM. n = 3-5 per experiment. Experiments are representative of at least 2 independent experiments. Statistical comparisons were performed with Student’s t test (C – E) or one-way ANOVA (A, B): p < 0.05; **p < 0.01; ***p < 0.001.

We also investigated whether broad depletion of the commensal microbiome, and thus SCFAs, was associated with alterations to Siglec-F^hi^ and Siglec-F^int^ lung eosinophils. To this end, we characterised lung eosinophils in specific pathogen free (SPF) mice compared to germ-free (GF) animals. GF mice had reduced frequencies of eosinophils in the BM (**Fig. 3 E**), whereas increased proportions were found in the lung (**Fig. 3 E and F**). Notably, Siglec-F^hi^ eosinophils that expressed CD101 were present in the lungs of non-allergic GF mice, which were barely detected in SPF animals (**Fig. 3 G**). Suggestive of augmented metabolic activation, the population of Siglec-F^hi^ eosinophils present in the lungs of GF animals also expressed CD98 (**Fig. 3 G**).

Overall, these findings demonstrate that commensal-derived factors, in particular butyrate, can modify the Siglec-F^hi^ eosinophil compartment in terms of frequency but also impact their activation state.

### The butyrate and niacin receptor GPR109A is selectively expressed by Siglec-F^hi^ eosinophils

Alongside its HDAC inhibitor activity, butyrate can also impact cell function via G protein-coupled receptors (GPCRs). Several GPCRs including GPR41, GPR43 and GPR109A, are reported to be receptors for butyrate (Kasubuchi et al., 2015; Dang and Marsland, 2019). We investigated whether Siglec-F^hi^ eosinophils expressed butyrate receptors during allergic-type inflammation. To this end, we first sorted Siglec-F^hi^ and Siglec-F^int^ eosinophils from the lung of HDM-treated mice and assessed their expression of butyrate-responsive GPCRs. Both Siglec-F^int^ and Siglec-F^hi^ lung eosinophils stained positive at low levels for GPR43 (**Supplementary Fig. 4 A**), while Siglec-F^int^ eosinophils most evidently expressed GPR41 (**Supplementary Fig. 4 B**). Notably, GPR109A was most apparent on Siglec-F^hi^ eosinophils suggesting a potential mechanism by which Siglec-F^hi^ eosinophils specifically could be impacted by SCFAs (**Fig. 4 A**).

**Figure 4:**
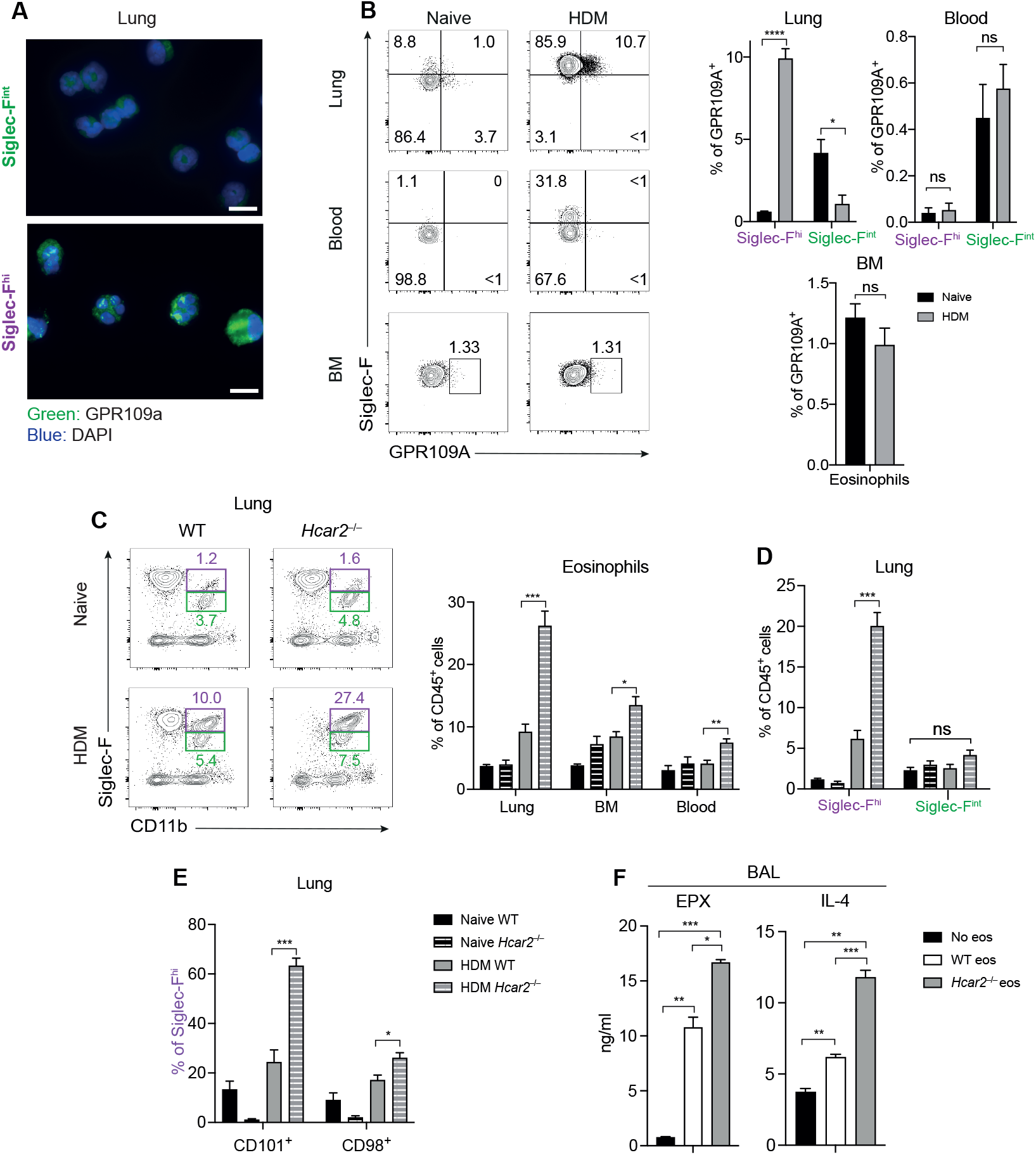
GPR109A-deficiency is associated with enhanced eosinophil activation. **(A)** Representative staining of sorted Siglec-F^int^ and Siglec-F^hi^ eosinophils from the lungs of HDM-treated mice for GPR109A (green) and counterstained with DAPI (blue) (scale bar 10 μm). **(B)** The percentage of eosinophils expressing GPR109A was quantified by flow cytometry in the lungs, BM and blood of naïve and HDM-treated mice. **(C)** Total eosinophils were quantified by flow cytometry in the lungs of naïve or HDM-treated WT and *Hcar2^−/−^* mice and expressed as a frequency of CD45^+^ cells. **(D)** Siglec-F^hi^ and Siglec-F^int^ eosinophils were quantified by flow cytometry in the lungs of naïve or HDM-treated WT and *Hcar2^−/−^* mice and expressed as a frequency of CD45^+^ cells. **(E)** The percentage of Siglec-F^hi^ eosinophils expressing CD101 and CD98 was quantified by flow cytometry in the lungs of naïve or HDM-treated WT and *Hcar2^−/−^* mice. **(F)** Levels of EPX and IL-4 were determined by ELISA from the BAL of HDM-treated ΔdblGATA mice that received intranasally transferred BM eosinophils from either WT or *Hcar2^−/−^* mice, or no eosinophils as a control. Error bars show ± SEM. n = 3-5 per experiment. Data are representative of at least 2 independent experiments. Statistical comparisons were performed with Student’s t test (B) or one-way ANOVA (C, D, E, F): p < 0.05; **p < 0.01; ***p < 0.001.

Flow cytometric analysis of Siglec-F^hi^ eosinophils also revealed that lung Siglec-F^hi^ eosinophils expressed increased levels of GPR109A compared to Siglec-F^int^ eosinophils in allergic mice (**Fig. 4 B**). GPR109A-staining was not observed on lung Siglec-F^hi^ eosinophils from *Hcar2^−/−^* (gene encoding GPR109A) animals (**Supplementary Fig. 4 C**), confirming the specificity of the GPR109A antibody. GPR109A-staining was also not detected on blood Siglec-F^hi^ eosinophils, blood Siglec-F^int^ eosinophils and BM eosinophils **(Fig. 4 B**), as was observed for CD98 and CD101 expression on blood and BM eosinophil subsets (**Fig. 2 E**).

Given that oral administration of the GPR109A-ligand butyrate, reduced the frequency and activation of Siglec-F^hi^ eosinophils in the lung during allergic-type inflammation (**Fig. 3 A – D**), we investigated whether mice lacking GPR109A (*Hcar2^−/−^*) would develop a reciprocal increase in activated Siglec-F^hi^ eosinophils in the same setting. While in naïve animals, eosinophils in the lung, blood and BM were unchanged between *Hcar2^−/−^* mice and WT controls, greater frequencies of eosinophils were evident in *Hcar2^−/−^* mice across all compartments following HDM-treatment (**Fig. 4 C**). In the lung, *Hcar2^−/−^* mice had an enhanced Siglec-F^hi^ eosinophil compartment with no alterations in Siglec-F^int^ eosinophils (**Fig. 4 D**). Siglec-F^hi^ eosinophils were more activated based on their expression of CD98 and CD101 (**Fig. 4 E**). Alterations to eosinophil populations in *Hcar2^−/−^* mice was associated with a globally augmented type 2 immune response, as evidenced by increased expression of transcripts for the Th2-cytokines *Il4* and *Il13* as well as the effector molecules *Retnla* (encoding RELM-α) and *Chil3* (encoding Ym1) (**Supplementary Fig. 4 D**).

To understand whether the impact of GPR109A was cell-intrinsic to eosinophils, we performed eosinophil transfer experiments. BM eosinophils from WT and *Hcar2^−/−^* mice were isolated and intranasally transferred into the lung environment of eosinophil-deficient (ΔdblGATA) mice during the challenge phase of HDM-mediated allergy. Assessment of EPX release in the BAL, as a measure of eosinophil activation, revealed that ΔdblGATA mice that received *Hcar2^−/−^* eosinophils released higher amounts of EPX and IL-4 compared to ΔdblGATA mice that received WT BM eosinophils (**Fig. 4 F**).

Taken together these findings reveal that the butyrate receptor GPR109A is an important receptor regulating allergic-type lung inflammation that has the potential to directly modulate the function of glycolytic Siglec-F^hi^ eosinophils. We next sought to understand the potential of GPR109A-ligands as candidate modulators of inflammation, preferentially targeting Siglec-F^hi^ lung eosinophils.

### Administration of GPR109A agonists suppresses allergic-type inflammation and the Siglec-F^hi^ eosinophil population

Butyrate is not the only agonist of GPR109A. β-hydroxybutyrate and niacin (also known as nicotinic acid or vitamin B3) can also bind and activate the receptor (Singh et al., 2014). β-hydroxybutyrate, like butyrate, has been reported to also act on cell function by inhibiting HDACs (Shimazu et al., 2013). Contrasting this, niacin does not have these histone-dependent modulatory effects that are independent of GPR109A (Lorenzen et al., 2001).

To investigate whether GPR109A agonism could be used to suppress allergic inflammation, at least in part through its effects on eosinophils, we intranasally treated animals with niacin at both the priming and challenge stages of the HDM model (**Supplementary Fig. 5 A**). Treatment led to a dramatic decrease in frequencies of eosinophils in the lung (**Fig. 5 A**) and more specifically impacted the Siglec-F^hi^ population, while the Siglec-F^int^ population was unaffected in terms of frequency (**Fig. 5 B**). This was also associated with alterations to the activation of the Siglec-F^hi^ population as both CD98 and p-mTOR were downregulated (**Fig. 5 B**). Additionally, as for oral butyrate treatment (**Supplementary Fig. 3 B**) gene expression of cytokines and effector molecules associated with type 2 immunity were also decreased (**Supplementary Fig. 5 B**). Of note, unlike oral administration of butyrate, intranasal niacin treatment had no impact on BM eosinophil frequency (**Fig. 5 A**).

**Figure 5:**
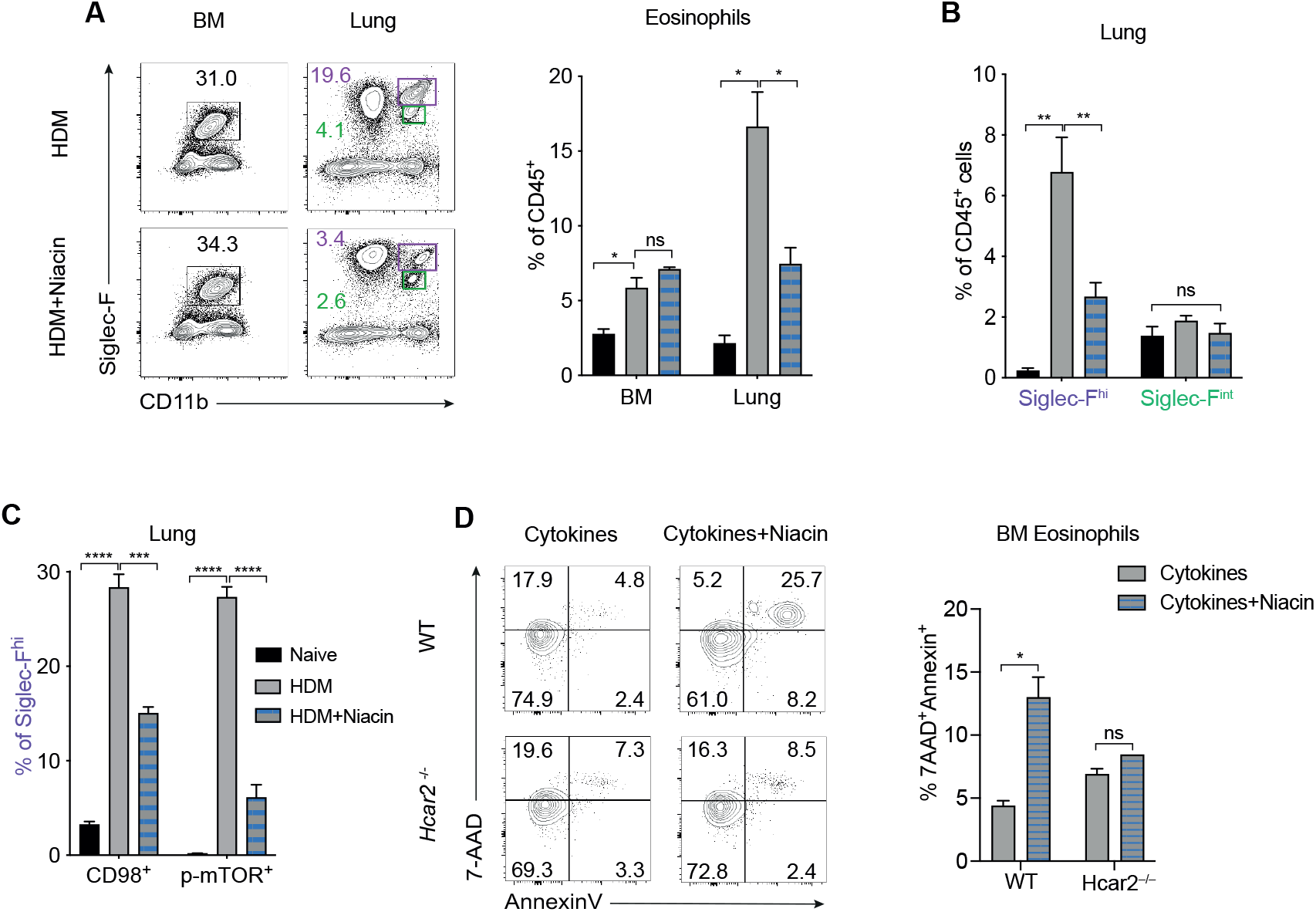
Treatment with niacin reduces eosinophil frequency and activation. **(A)** Eosinophils were quantified by flow cytometry in the lungs and BM of naïve mice and mice treated with HDM alone or with niacin and expressed as a percentage of CD45^+^ cells. **(B)** Siglec-F^hi^ and Siglec-F^int^ eosinophils were quantified by flow cytometry in the lungs of naïve mice and mice treated with HDM alone or with niacin and expressed as a percentage of CD45^+^ cells. **(C)** The percentage of Siglec-F^hi^ eosinophils expressing CD98 and p-mTOR was quantified by flow cytometry in the lungs of naïve mice and mice treated with HDM alone or with niacin. **(D)** The percentage of BM eosinophils from WT and *Hcar2^−/−^* mice expressing Annexin V and 7-AAD was quantified by flow cytometry following culture with IL-33, GM-CSF and IL-5 in the presence or absence of niacin for 24 hours. Error bars show ± SEM. n = 3-5 per experiment. Data are representative of 3 independent experiments. Error bars show ± SEM. Statistical comparisons were performed with Student’s t test (C) one-way ANOVA (A, B): p < 0.05; **p < 0.01; ***p < 0.001.

For neutrophils, GPR109A signaling is associated with apoptosis via inhibition of HDACs (Kostylina et al., 2008). We cultured mixed BM cells in the presence of eosinophil survival factors found in the inflamed lung environment (IL-5, IL-33 and GM-CSF) alone or with niacin and assessed eosinophil apoptosis. Niacin increased apoptosis and necrosis in WT eosinophils compared to stimulation with the lung cytokines alone (**Fig. 5 D**). This effect was abrogated in *Hcar2^−/−^* BM eosinophils, suggesting that one mechanism by which GPR109A signals may modulate the Siglec-F^hi^ population is by inducing their apoptosis (**Fig. 5 D**).

Taken together, our findings demonstrate that Siglec-F^hi^ eosinophils are responsive to signals through the GPR109A receptor. This presents a new mechanism that could be used to target allergic inflammation. Therapeutic administration of GPR109A ligands can be used as a strategy to selectively remove inflammatory eosinophils, likely via apoptosis, while sparing regulatory/homeostatic populations that are increasingly understood to be important for normal physiological function and immunoregulation of organs.

## DISCUSSION

Here we have shown that Siglec-F^hi^ eosinophils and Siglec-F^int^ eosinophils represent two phenotypically and metabolically distinct populations of eosinophils found in the airways of allergic mice. Siglec-F^hi^ eosinophils exhibited a pro-inflammatory phenotype and function that were correlated with higher glycolytic activity and activation of the mTOR signaling pathway. Additionally, we identified Siglec-F^hi^ eosinophils expressing CD98 and CD101 as a common feature of lung inflammation. Most importantly, we demonstrated that lung Siglec-F^hi^ eosinophils express GPR109A and that agonism of this receptor can directly affect eosinophil phenotype and function. Therapeutic administration of the GPR109A agonists niacin and butyrate resulted in inhibition of allergic-type inflammation and reduced Siglec-F^hi^ eosinophil frequencies and activation. Thus, GPR109A may provide a powerful new target to modulate Siglec-F^hi^ inflammatory eosinophils during allergic disease, while sparing Siglec-F^int^ eosinophils with regulatory or homeostatic functions.

Although little is known about eosinophil metabolism, our findings are broadly in agreement with published data showing that human blood eosinophils are plastic cells able to upregulate glycolysis and the tricarboxylic acid (TCA) cycle upon activation (Jones et al., 2020). Moreover, it has been suggested that, although both human eosinophils and neutrophils use glycolysis, eosinophils also access additional pathways such as glucose oxidation and mitochondrial oxidative phosphorylation (Porter et al., 2018). This supports the idea that eosinophils show significant metabolic flexibility, which allows them to better adapt to tissue requirements. In particular, we identified that Siglec-F^hi^ lung eosinophils were more glycolytic than the Siglec-F^int^ lung eosinophil population. To our knowledge this is the first demonstration of differential metabolism between distinct populations of eosinophils *in vivo*. Inflamed tissues are often hypoxic, due to increased cellular oxygen demand and reduced oxygen availability. Hypoxia is a pro-inflammatory stimulus, which activates HIF-1α signaling, resulting in a metabolic shift from oxidative phosphorylation to glycolysis. Enhanced glycolysis in eosinophils may therefore allow them to perform their functions in environments with limited oxygen availability, such as the allergic lung (Porter et al., 2018).

Previous studies documented the existence of two separate populations of eosinophils in murine models of allergic asthma (Mesnil et al., 2016; Abdala Valencia et al., 2015). Siglec-F^int^ eosinophils were reported not to be affected by allergen exposure and to have regulatory functions, whereas Siglec-F^hi^ eosinophils were shown to increase in response to allergen inhalation and to be pro-inflammatory (Mesnil et al., 2016). Similarly, our results showed a simultaneous decrease in the levels of Siglec-F^int^ eosinophils with an increase in Siglec-F^hi^ eosinophils following HDM administration. Phenotypically, Siglec-F^hi^ eosinophils, but not Siglec-F^int^ eosinophils, have been reported to express CD101 (Mesnil et al., 2016), which we confirmed in our analysis. In addition, we saw that Siglec-F^hi^ eosinophils expressed elevated levels of the amino acid transporter CD98. CD98 is known to support cell growth and survival by multiple mechanisms; it allows amino acid transport by coupling with LAT1 and its association with integrins is linked to regulation of integrin function (Cormerais et al., 2016). Our data showed that CD98 expression by Siglec-F^hi^ eosinophils correlated with concurrent activation of the mTOR signaling pathway, as well as increased glycolytic activity.

Cellular metabolism is essential in determining immune cell function and metabolic changes are crucial in dictating how cells respond to a variety of signals (Ganeshan et al., 2014). Our data show that Siglec-F^hi^ eosinophils are more pro-inflammatory and responsive to stimulation than Siglec-F^int^ eosinophils, as they released higher amounts of IL-4 and EPX *in vitro*. These data further support the concept that the Siglec-F^hi^ subset of eosinophils are highly tailored to alterations in the tissue microenvironment and that this involves their metabolic reprogramming. Of note, acquisition of these features likely only occurs in the lung as BM and blood circulating eosinophils did not express CD101 or CD98.

Although we and others have pointed out changes in eosinophil phenotype in allergic asthma, it is not known whether similarities or differences in eosinophils exist across murine models of lung inflammation., In the settings of type 2 parasite-induced lung inflammation and type I IFN favouring influenza infection, we detected Siglec-F^hi^ eosinophils and Siglec-F^int^ eosinophils in the lungs, with Siglec-F^hi^ eosinophils expressing CD98 and CD101. These data reveal that Siglec-F^hi^ eosinophils with an activated phenotype are a common characteristic of lung inflammatory responses.

Interestingly, our phenotypic analysis of lung eosinophils suggested that Siglec-F^hi^ eosinophils expressed higher levels of the butyrate receptor GPR109A (Macia et al., 2015) compared to Siglec-F^int^ eosinophils. GPR109A is known to be expressed in adipocytes and immune cells, including macrophages, DCs and neutrophils (Tunaru et al., 2003; Singh et al., 2015; Zandi-Nejad et al., 2013) but not previously by murine eosinophils. In humans, mature neutrophils but not eosinophils have been reported to express GPR109A (Kostylina et al. 2008). However, this study was based on analysis of human blood rather than cells from tissues, such as the lung. Our data and others (Weller and Spencer, 2017; Olbrich et al., 2020) have shown that eosinophils in the lung microenvironment are phenotypically and functionally distinct from those in blood or other tissues such as the gut, suggesting that the properties of circulatory eosinophils are not representative of eosinophils from different tissue compartments. Thus, identification of new therapeutic strategies to locally deplete pro-inflammatory eosinophils maybe dependent on studying eosinophils in the inflamed organ.

Several studies suggest that SCFAs, such as butyrate, have beneficial effects on inflammatory disease pathogenesis, including allergic asthma (Maslowski et al., 2009; Trompette et al, 2014; Cait et al., 2018). Although butyrate has recently been shown to affect eosinophil survival and migration (Theiler et al., 2019), it is still unclear whether lung eosinophils can be directly targeted by SCFAs during allergic lung inflammation. We found that local agonism of the butyrate receptor GPR109A reduced eosinophil activation and frequency demonstrating it can be locally targeted. *In vitro* stimulation of GPR109A suggested that one direct mechanism by which GPR109A-signaling impacted eosinophils was by faciliating apoptosis, a process similarly reported in neutrophils (Kostylina et al., 2008).

The identification of different asthma phenotypes has led to development of more tailored treatments, such as biologic therapies to target specific inflammatory mediators (Brussino et al., 2018). Considering the fundamental role of IL-5 in eosinophil differentiation and survival, anti-IL-5 and anti-IL-5Rα antibodies have been developed (Wechsler et al., 2012). However, although treatment with these therapeutics reduces circulating eosinophils, they do not provide a significant clinical improvement in some asthma patients (Borish, 2016; Brussino et al., 2018). With this work we provide evidence for alternative targatable pathways that could be agonised to prevent or reduce eosinophil activation, including the CD98-mTOR signaling pathway and glycolysis. We also highlight the importance of the gut microbiota and derived metabolites in influencing peripheral immunity and show that ligation of GPR109A by niacin-like ligands may be able to specifically target Siglec-F^hi^ eosinophils and counteract inflammatory responses in allergic lung inflammation.

This study indicates that lung eosinophils are plastic cells that can adapt to tissue requirements by changing their phenotype, functional characteristics and metabolism during type 2 lung inflammatory responses. In addition, we demonstrate GPR109A agonism as a means to regulate pro-inflammatory eosinophils during type 2 inflammation and, thus, a candidate therapeutic intervention to slow the progression of inflammatory lung disease.

## Acknowledgements

We thank: Prof. Mark Travis for providing the PR8 strain of influenza; Prof. Andrew MacDonald for providing *Schistosoma Mansoni*; Prof. Judith Allen for providing *Nippostrongylus brasiliensis*; and Joana Costa Oliveira and Sabrina Tamburrano for careful reading of the manuscript. *Hcar2^−/−^* mice were kindly provided by Prof. Stefan Offermanns (Max-Planck-Institute for Heart and Lung Research, Bad Nauheim) and Prof. Markus Schwaninger (University of Lübeck). We thank Dr. Tara Sutherland and Dr. Soren Bienke for helpful discussion. We acknowledge the support of the Bioimaging, Flow Cytometry and Biological Services core facilities at The University of Manchester. The University of Manchester Axenic and Gnotobiotic facility used was established with the support of the Wellcome Trust (097820/Z/11/B).

This work was supported by a Kennedy Trust for Rheumatology Research Senior Fellowship awarded to J. R. Grainger and a grant awarded to the Manchester Collaborative Centre for Inflammation Research (MCCIR) (funded by a precompetitive open innovation award from GlaxoSmithKline).

