## Supplementary Figures for "Distinct eosinophil subsets are modulated by agonists of the commensal-metabolite and vitamin B3 receptor GPR109A during allergic-type inflammation"

**A**

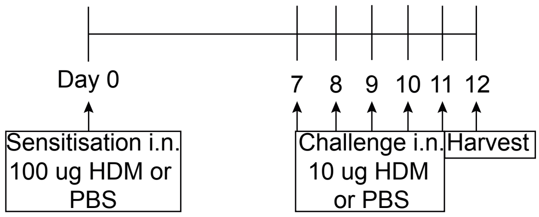


**B**

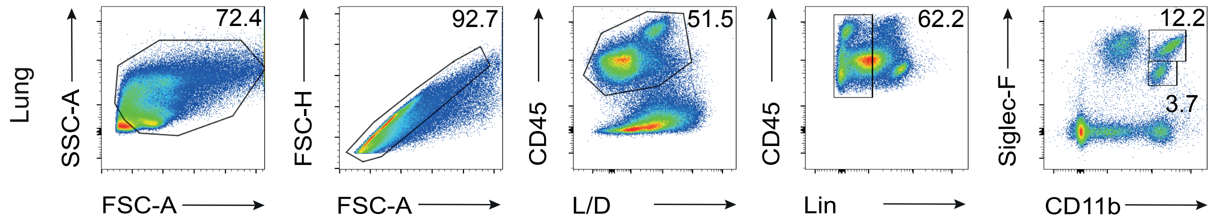


**Supplementary Figure 1**

**(A)** Experimental timeline. Female WT BALB/c mice (8-12 weeks) were sensitised intranasally (i.n.) on day 0 with 100 µg HDM and challenged i.n. on days 7-11 with 10 µg HDM. Control mice were treated i.n. at the same time points with PBS. Mice were euthanised 24 hours after the last challenge (on day 12). **(B)** Gating strategy to identify lung eosinophils. Cells were identified as follows: SSC-A versus FSC-A to remove cellular debris; FSC-H versus FSC-A for doublet exclusion; Live-Dead (L/D) versus CD45 to identify live haematopoietic populations; lineage (Lin) versus CD45 to exclude contaminating populations (lineage defined as TCRβ, B220, NK1.1, Ly6C and Ly6G to exclude T cells, B cells, NK cells, monocytes/macrophages and neutrophils, respectively); Siglec-F versus CD11b to identify eosinophils ($CD11b^{+}Siglec\text{-}F^{hi/int}$) and exclude alveolar macrophages (AMs) ($Siglec\text{-}F^{+}CD11b^{-}$).

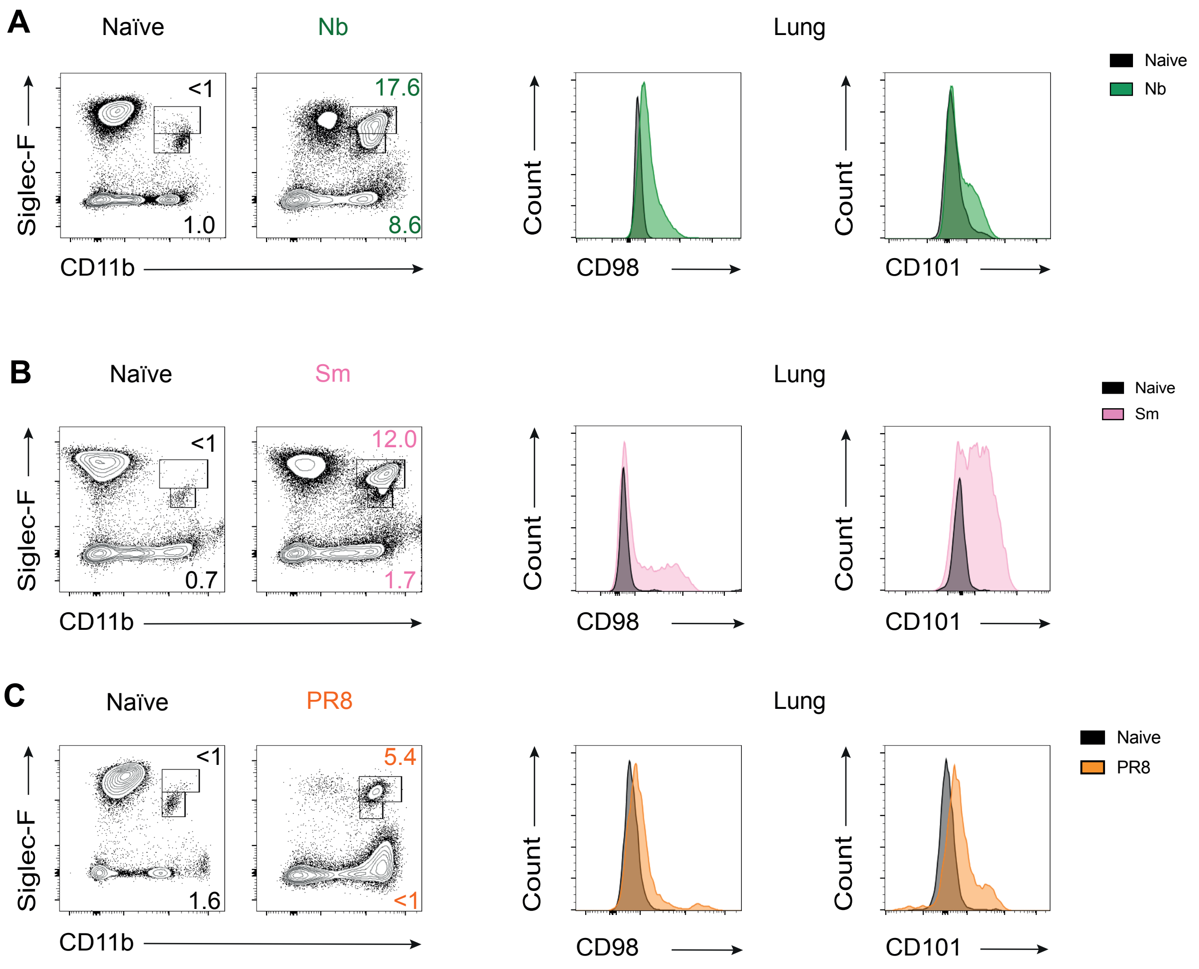


**Supplementary Figure 2**
**(A – C)** Representative flow cytometry plots showing lung eosinophils from naïve and mice infected with (A) N. brasiliensis, **(B)** S. Mansoni and **(C)** PR8. Histograms showing expression of CD98 and CD101 by lung eosinophils from naïve and mice infected with *N. brasiliensis*, *S. mansoni* and PR8.

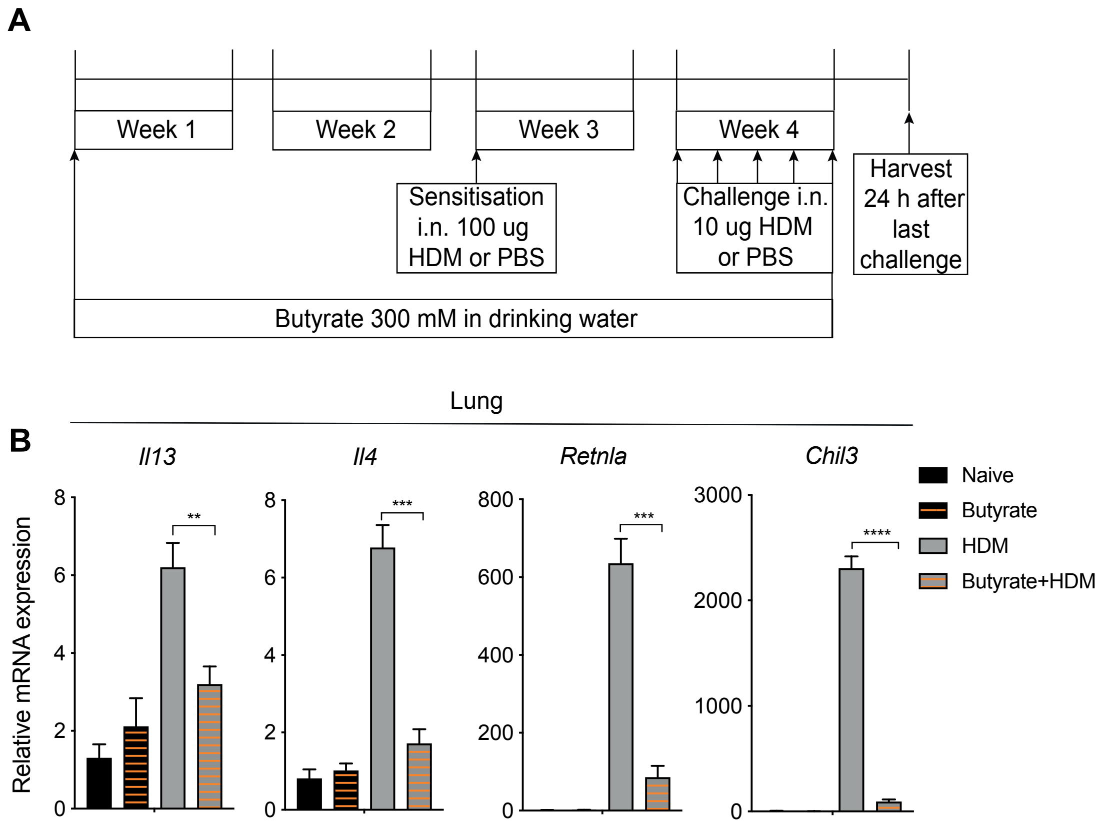


**Supplementary Figure 3**

**(A)** Experimental timeline for butyrate treatment during HDM. Female WT BALB/c mice, 8-12 weeks, were administered 300 mM butyrate in the drinking water for two weeks prior and then during HDM treatment. Mice were euthanised 24 hours after the last challenge with HDM (on day 12). **(B)** Gene expression of *Il4*, *Il13*, *Chil3* (encoding Ym-1) and *Retnla* (encoding Relm-α) was assessed in homogenised lung tissue of naïve mice, mice treated with butyrate alone, mice treated with HDM alone and mice treated with HDM and butyrate. Error bars show ± SEM. n = 3-5. Data are representative of at least 3 independent experiments. Statistical comparisons were performed with one-way ANOVA: $p < 0.05$; $^{**}p < 0.01$; $^{***}p < 0.001$.

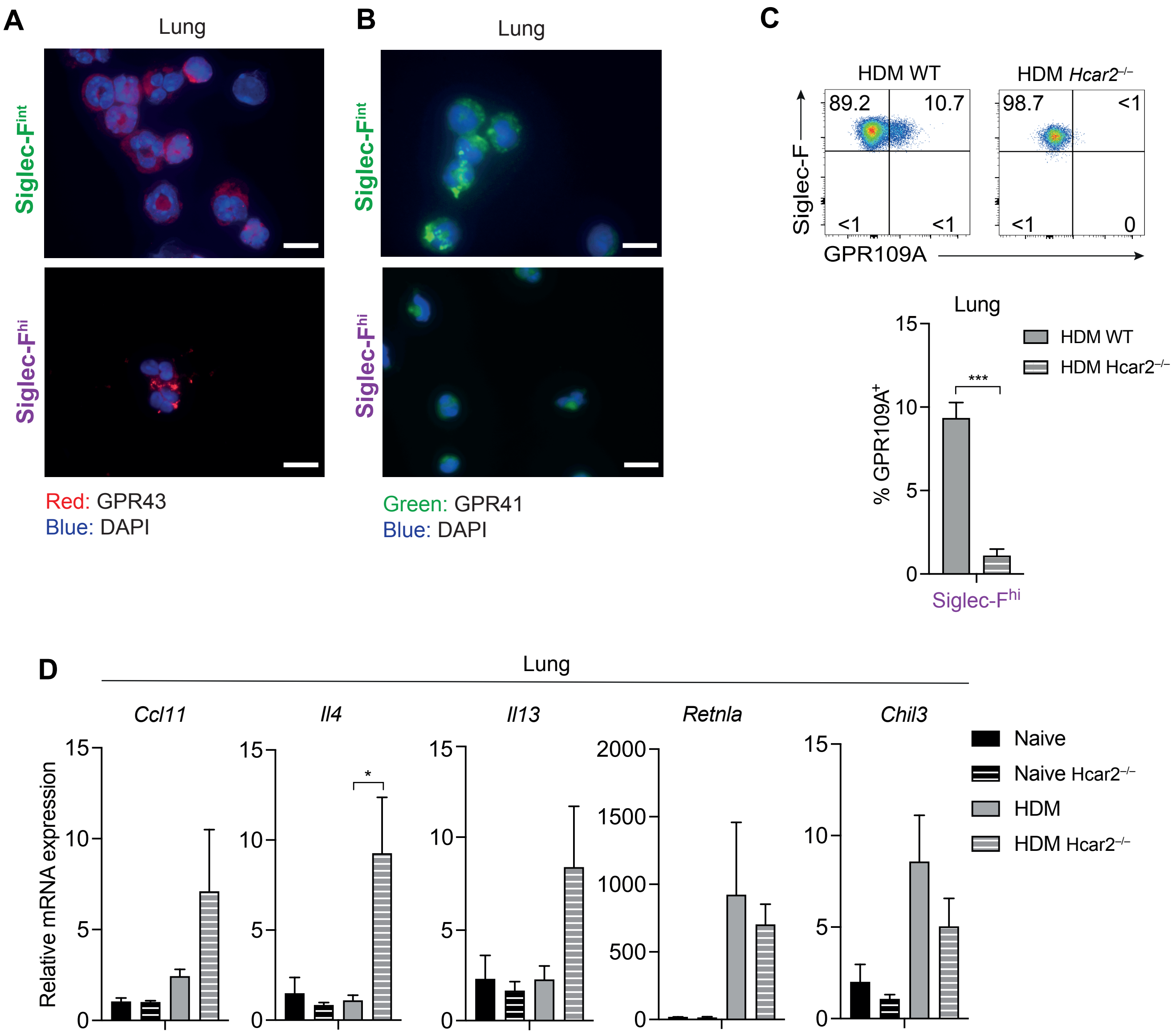


**Supplementary Figure 4**

**(A)** Representative staining of sorted Siglec-F$^{int}$ and Siglec-F$^{hi}$ eosinophils from the lungs of HDM-treated mice for GPR43 (red) and counterstained with DAPI (blue) (scale bar 10 μm). **(B)** Representative staining of sorted Siglec-F$^{int}$ and Siglec-F$^{hi}$ eosinophils from the lungs of HDM-treated mice for GPR41 (green) and counterstained with DAPI (blue) (scale bar 10 μm). **(C)** The percentage of Siglec-F$^{hi}$ eosinophils expressing GPR109A was quantified by flow cytometry in the lungs of HDM-treated WT and *Hcar2*$^{-/-}$ mice. **(D)** Gene expression of *Ccl11*, *Il4*, *Il13*, *Chil3* and *Retnla* was established in lung tissue of naïve or HDM-treated WT and *Hcar2*$^{-/-}$ mice. Error bars show ± SEM. n = 3-5 per experiment. Data are representative of at least 2 independent experiments. Statistical comparisons were performed with Student's t test (C) or one-way ANOVA (D): $p < 0.05$; **$p < 0.01$; ***$p < 0.001$.

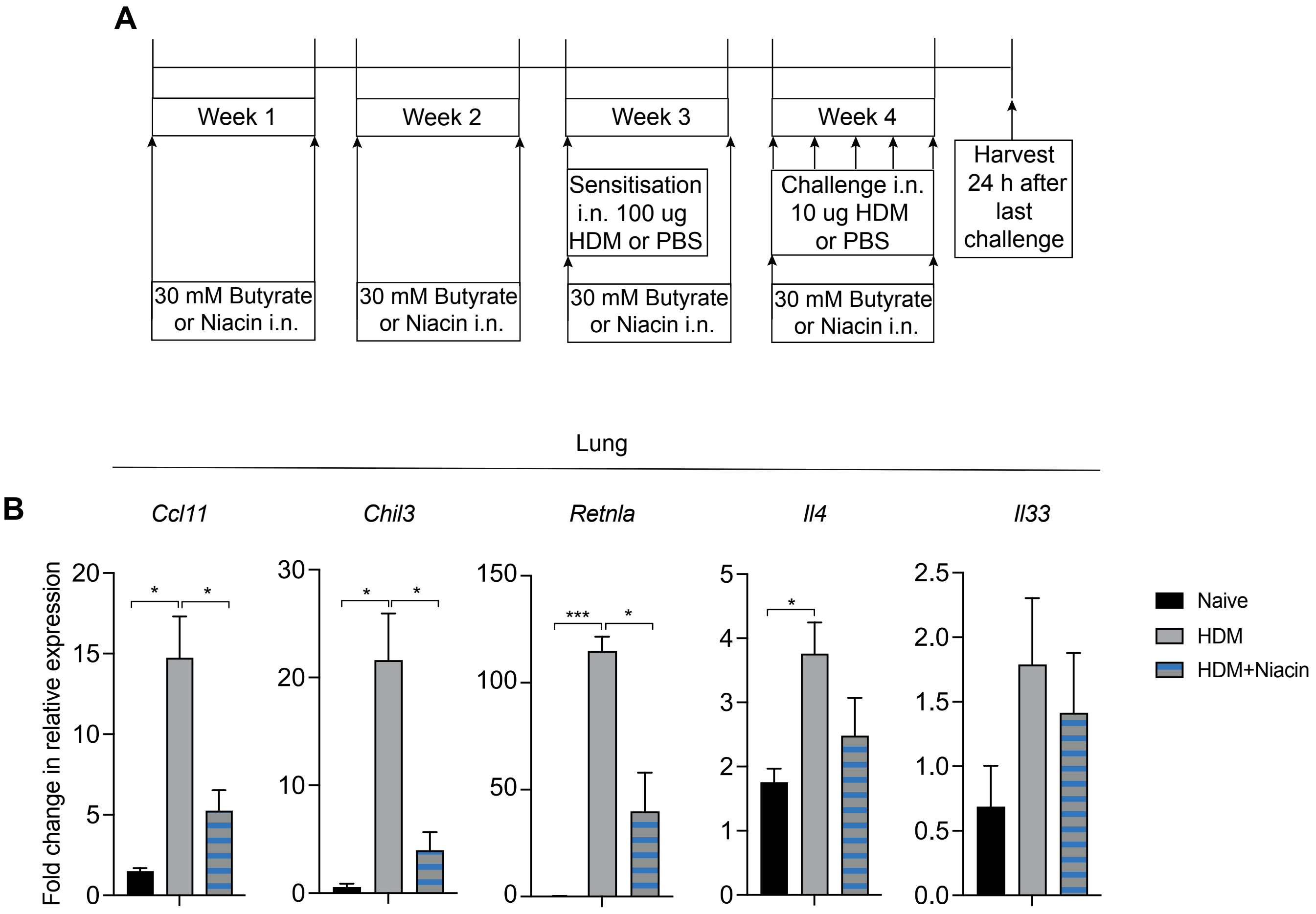


**Supplementary Figure 5**

**(A)** Experimental timeline. Female BALB/c WT mice, 8-12 weeks old, were administered 30 mM niacin I.N. twice a week for two weeks prior and during HDM treatment. Mice were euthanised 24 hours after the last challenge with HDM (on day 12). **(B)** Gene expression of *Ccl11*, *Il4*, *Il33*, *Chil3* and *Retnla* in lung tissue of naïve mice and mice treated with HDM alone or with niacin. Data are representative of 3 independent experiments. n = 3-5. Error bars show ± SEM. Statistical comparisons were performed with one-way ANOVA: $p < 0.05$; $^{**}p < 0.01$; $^{***}p < 0.001$.
